# A system for gene expression noise control in yeast

**DOI:** 10.1101/359042

**Authors:** Max Mundt, Alexander Anders, Seán Murray, Victor Sourjik

**Author notes:** Correspondence to: Victor Sourjik.

## Abstract

Gene expression noise arises from stochastic variation in the synthesis and degradation of mRNA and protein molecules and creates differences in protein numbers across populations of genetically identical cells. Such variability can lead to imprecision and reduced performance of both native and synthetic networks. In principle, gene expression noise can be controlled through the rates of transcription, translation and degradation, such that different combinations of those rates lead to the same protein concentrations but at different noise levels. Here, we present a “noise tuner” which allows orthogonal control over the transcription and the mRNA degradation rates by two different inducer molecules. Combining experiments with theoretical analysis, we show that in this system the noise is largely determined by the transcription rate whereas mean expression can be independently adjusted by mRNA stability. This noise tuner enables twofold changes in gene expression noise over a fivefold range of mean protein levels. We demonstrated the efficacy of the noise tuner in a complex regulatory network by varying gene expression noise in the mating pathway of *Saccharomyces cerevisiae*, which allowed us to control the output noise and the mutual information transduced through the pathway. The noise tuner thus represents an effective tool of gene expression noise control, both to interrogate noise sensitivity of natural networks and enhance performance of synthetic circuits.

## Introduction

All processes in cells are subject to variation in the numbers or states of involved molecules. These random fluctuations cause gene expression noise, which is the variation of the numbers of gene products across a population of clonally identical cells. Experimentally one can distinguish between two types of noise that affect expression of a particular gene (*1*). Extrinsic or global noise is caused by the fluctuations of concentrations, state, or location of factors that affect multiple genes or even the entire cell, such as the number of ribosomes as a limiting factor for overall protein production (*2*). Intrinsic or gene‐specific noise is caused by the stochasticity of the biochemical reactions during each step of the expression of an individual gene, including transcriptional (e.g. transcription initiation), post‐transcriptional (e.g. mRNA degradation), (co‐)translational (e.g. translation initiation) and post‐translational processes (e.g. protein degradation) (*3*).

While in some cases variability in expression within a population of cells might be beneficial, as in the case of environmental stress response (*4*), expression noise typically reduces cellular fitness (*5*) and is generally counter‐selected (*6*). Low noise is especially important for cellular processes that require a certain stoichiometry of involved proteins (*7, 8*). Similarly, noise control is important to achieve reliable function of complex synthetic genetic circuits (*9*).

Consequently, several strategies for noise reduction have been recognized in living organisms and applied in the design of synthetic genetic circuits. One common strategy is to use negative feedback loops, e.g. by transcriptional repression (*10*). However, such negative feedback control of gene expression has been shown to have very limited efficiency and comes at high cost (*11*). Decoupling noise from the mean expression has also been achieved by altering the expression of upstream regulators of the target gene (e.g. *12, 13*) or by inserting inducible promoters with different noise characteristics (*14*).

In principle, noise can be effectively controlled via the basic rate constants of gene expression (*15*). In the simplest model of gene expression, the steady state concentration of a protein is governed by the rate constants of four processes: transcription, mRNA degradation, translation, and protein degradation. Here, the same amount of a protein can be produced when the gene is transcribed at a high rate but transcripts have a low translation rate, or when the gene is transcribed at a low rate but the translation rate is high. In the first case, many mRNA molecules are translated rarely, whereas in the second case only few transcripts are made but translated multiple times. The first scenario generally results in lower variation of protein concentration over time, because stochasticity of transcription is averaged out due to a high number of transcripts (*16*).

Here, we describe an approach to decouple the mean expression from the expression noise of a gene by using two different inducers, independently controlling the transcription rate and the mRNA degradation rate, respectively. Stochastic simulations of an analytical model that links gene expression to population‐level distributions of protein showed that the noise in this system is mainly governed by transcriptional bursts at low promoter activities. Finally, we used our noise tuner to control signaling noise and therefore the amount of conveyed information in the yeast mating pathway, a prototypic signaling pathway for synthetic biology (e.g. *17, 18, 19*).

## Results and discussion

### Design and implementation of noise tuner

In order to decouple mean gene expression levels from expression noise in *Saccharomyces cerevisiae*, we built a system that allows independent control of both, transcription rate and mRNA degradation rate for a specific gene. This system that we term “noise tuner” employs a doxycycline‐inducible Tet‐promoter to vary mRNA production and a ribozyme regulated by a small molecule theophylline (*20*) to adjust the mRNA half‐life (Figure 1a). Doxycycline binds to the constitutively expressed reverse tetracycline trans‐activator (rtTA, *21*), which is then recruited to Tet‐operator (TetO) sites within the Tet‐promoter and initiates transcription. The mRNA degradation rate is controlled by a ribozyme sequence in its 3’ untranslated region (3’‐UTR). This sequence folds into a stem loop and exhibits autocatalytic endoribonuclease activity that is inhibited by binding of theophylline, which therefore increases gene expression (Figure S1). Additionally, a short synthetic transcriptional terminator (*22*) is placed downstream of the gene. Hereinafter, we will refer to a combination of 3’‐UTR and transcriptional terminator as a 3’ regulatory region (3’‐RR).

**Figure 1.**
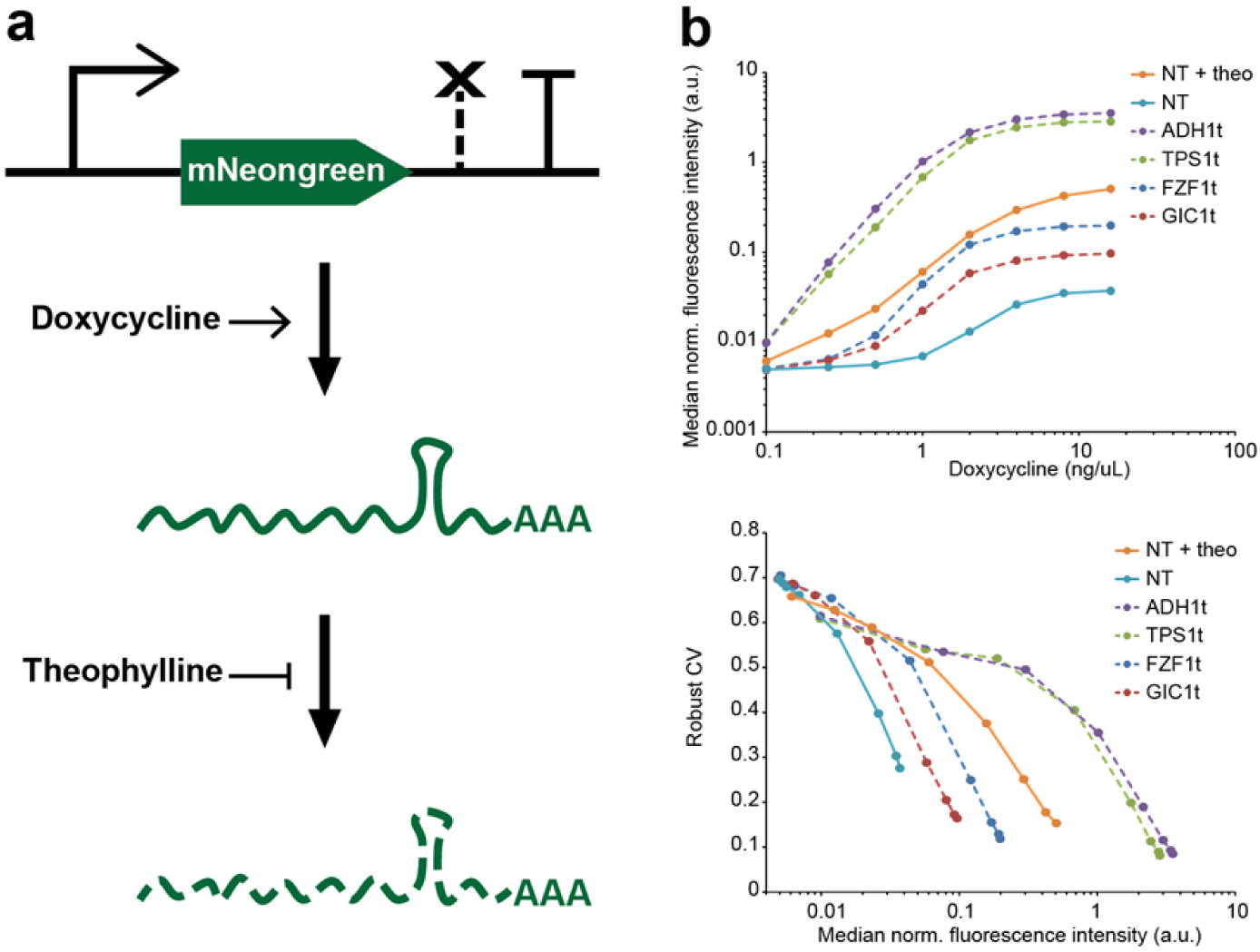
Design and benchmarking of a noise tuning system. **(a) Schematic depiction of the noise tuner.** In a Tet‐ON system *(21)* doxycycline activates transcription of the fluorescent mNeongreen reporter gene by binding to the constitutively expressed reverse tetracycline trans‐activator (rtTA, not shown). Addition of theophylline prevents mRNA degradation by disrupting the ribozyme cleavage activity in the 3’‐UTR. **(b) Comparison of the noise tuner with constructs harboring native terminator regions measured by flow cytometry.** All constructs are identical in their Tet promoter and mNeongreen fluorescent reporter gene and differ only in their terminator regions that confer low (FZF1t, GIC1t) or high (TPS1t, ADH1t) expression *(24)*. Top panel: Dose‐responses with 0 to 12.8 ng/μl doxycycline. The noise‐tuner transcript was either unstable (0 mM theophylline, “NT”) or fully stabilized (12.8 mM theophylline, “NT + theo”). Bottom panel: The noise tuner exhibits similar inverse noise‐median correlation as native terminators. Noise is given as the robust coefficient of variation (robust CV, see Methods). Median fluorescence intensities normalized to a constitutively expressed mTurquoise2 reporter gene are given in arbitrary units (a.u.). Higher median fluorescence intensities correspond to induction with higher doxycycline concentrations.

For proof of concept, we genomically integrated noise tuner modules controlling the expression of the yellow‐green fluorescent protein mNeongreen (*23*) into yeast and measured fluorescence in single cells by flow cytometry. Additionally, our reporter strains constitutively expressed a blue fluorescent protein mTurquoise2, which allowed for cell‐by‐cell normalization of the mNeongreen signal and thus effective elimination of extrinsic noise sources (see below).

First, we tested the efficacy of expression regulation for two different ribozymes, (L2b8‐47 and L2b8‐a1‐ t41, here termed Ribo 1 and Ribo 2; *20*), which yielded different absolute levels of protein expression but showed similar, approximately 13‐fold, relative changes upon addition of theophylline (Figure S1). If not mentioned otherwise, subsequent results reported here were obtained with ribozyme Ribo 2, which generally confers lower expression.

Next, we tested whether the synthetic 3’‐RR conferred expression levels within the range of native yeast 3’‐RRs. For this purpose, we compared the noise tuner containing Ribo 2 to constructs with the identical Tet‐promoter and mNeongreen sequences, but different native yeast 3’‐RRs, which have been shown to result in either very high or low reporter expression, respectively (*24*). We found that the noise tuner operates within the lower range of expression strengths (Figure 1b, upper panel); however, if required, higher overall expression can be achieved by employing the 3’‐RR with ribozyme Ribo 1 (Figure S1). We also calculated gene expression noise in these experiments. Here, the noise in protein levels, i.e., fluorescence intensities, across the population is given as the robust coefficient of variation (see Methods). We found that gene expression noise at a given expression intensity correlates with the overall expression strength conferred by a 3’‐RR (Figure 1b, lower panel).

We noticed that stronger transcriptional stimulation (i.e., doxycycline concentration) led to lower gene expression noise at a given median expression level. To further explore the relationship between transcription, mRNA degradation and noise, we adjusted steady state expression of the noise tuner with different combinations of doxycycline and theophylline concentrations to differentially control transcription and mRNA degradation rates. From that, we derived a comprehensive expression landscape (Figure S2) and a noise landscape which integrates the median expression of the fluorescence marker in a population of cells with the variation around that median (Figure 2a). In the noise matrix, “iso‐median” lines indicate same median expressions that have been obtained with different combinations of transcription and mRNA degradation rates (Figure 2a). We observed that while the noise levels at a given median expression were reduced with increasing transcription rate, noise was not affected by changes in mRNA degradation rate. As a consequence, when a given median expression level is reached using high transcription rate and low transcript stability, the population of cells exhibits a significantly (up to twofold) lower coefficient of variation as compared to populations with lower transcription rate and higher transcript half‐life (Figure 2b). Similar results with overall higher expression levels were obtained when using Ribo 1 for noise tuning (Figure S3a), whereas no effect of theophylline was observed for a control strain without the ribozyme (Figure S3b).

**Figure 2.**
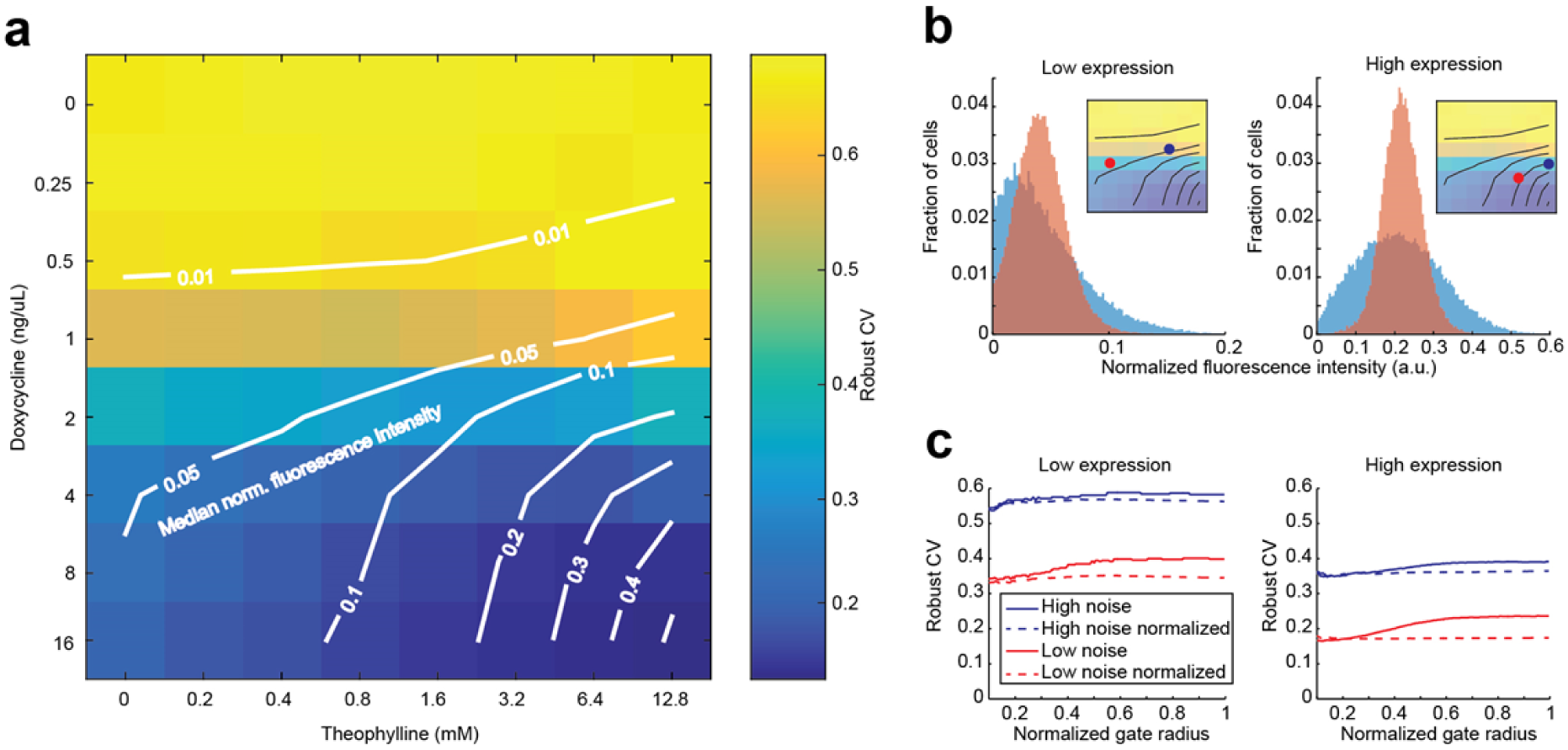
Gene expression noise is decoupled from mean expression by orthogonal control of transcription rate and mRNA degradation rate. **(a) Median‐noise landscape for noise‐tuner expression at different combinations of doxycycline and theophylline concentrations.** Colors indicate noise; lines indicate identical median normalized fluorescence, interpolated from the measured data (“iso‐median lines”). At intermediate expression range, noise levels can be adjusted for a given median expression by use of different combinations of doxycycline and theophylline. The measured robust CV decreases when a median is reached with higher transcription rates and correspondingly lower mRNA degradation rates. **(b) Example histograms of populations with similar median expression but different noise settings.** Populations with higher mNeongreen transcription and mRNA degradation rates (red) display lower heterogeneity than populations with lower rates (blue). Insets indicate the compared populations from **a. (c) Deconvolution of measured noise.** For populations shown in **b**, noise is plotted against the radius of a circular gate around the median forward (FSC) and side scatter (SSC) values. Decreasing gate sizes lead to more homogeneous FCS‐SSC populations, effectively filtering out the extrinsic component of the observed noise *(4)*. In contrast to autofluorescence‐subtracted raw data (solid lines), noise of autofluorescence‐subtracted data normalized to a constitutively expressed mTurquoise2 reporter (dashed lines) is virtually independent of the gate size.

Since the noise tuner adjusts gene‐specific factors of expression, it is expected to primarily control intrinsic noise, whereas global expression noise should be largely removed by normalization to mTurquoise2 levels. To verify this, we performed a reduced gate size analysis, which can be used in yeast to discriminate between extrinsic and intrinsic contributions to the observed noise (*4*). The analysis was done by selecting subsets of cells according to their forward (FSC) and side scatter (SSC) signals in flow cytometric measurements. FSC and SSC provide information about cell size and granularity, so that by reducing the subset size by applying a smaller gate, sources of extrinsic variation are filtered out. We found that noise for non‐normalized data was somewhat reduced for smaller, morphologically more similar subsets of cells, while it remained virtually unchanged for normalized data (Figure 2c). We conclude that extrinsic factors make only minor contribution to gene expression noise in our non‐normalized data and cell‐wise normalization to the constitutive reporter essentially eliminates those factors. Noise in the populations on different ends of the same iso‐median line changes up to twofold.

### Model of expression noise

To further examine the mechanisms behind different dependencies of noise tuner mean expression and noise on doxycycline and theophylline levels, we turned to a theoretical analysis of the distribution of protein copy numbers in a population of cells (*25*). In this model, the distribution of protein levels depends on only two parameters: 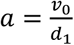, where *v*_0_ is the mRNA transcription rate and *d*_1_ is the protein degradation rate; and 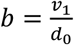, where *v*_1_ is the translation rate and *d*_0_ is the mRNA degradation rate (Figure 3a). The parameter *b* corresponds to the translational burst size – the average number of proteins translated per mRNA – while *a* can be interpreted as the average number mRNA transcribed during the lifetime of a single protein.

**Figure 3.**
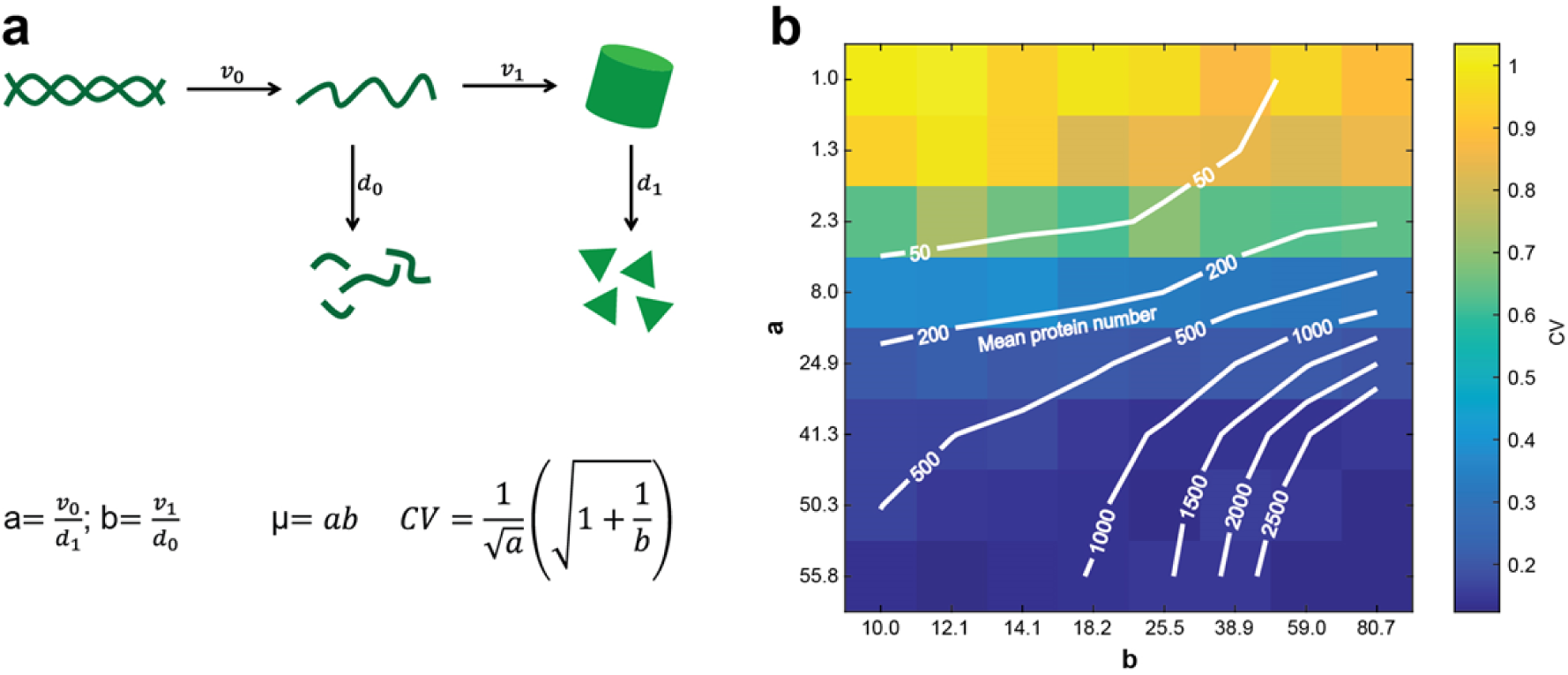
A simple mathematical model can reproduce the observed median‐noise relationship. **(a) Schematic depiction of the employed model.** The synthesis and degradation rates of mRNA and protein (v_0_ and d_0_ and v_1_ and d_1_, respectively) are used to calculate the coefficients a and b. The mean protein number μ and its noise (given as the CV) can be defined as a function of a and b *(25)*. **(b) Simulated protein number and CV as function of a and b.** Lines indicate same mean protein number, colors indicate CV. Scaling of a and b axes was chosen to correspond to experimentally observed expression intensities with different doxycycline and theophylline concentrations.

We note that if the burst size *b* ≫ 1 then the CV depends only on the parameter *a*, i.e. on the translation rate and the mRNA degradation rate. Large *b* implies that many proteins are produced from few mRNAs, which is indeed typically the case (see e.g. *26* and *27*). This independence of the CV on the burst size was further confirmed by stochastic simulations, which could reproduce the experimental observations in a biologically reasonable parameter space (see Methods). While the mean protein number was maximized when both, *a* and *b*, were high, the CV scaled with *a* and only weakly with *b* (Figure 3b).

### Tuning of information flow through a signaling pathway

Finally, we tested the operation of the noise tuner in the context of a cellular network to investigate whether changing the noise levels of individual network components could change the noise of the pathway output. We chose to probe noise sensitivity of the yeast mating pathway (Figure 4a), a prototypic MAPK pathway and a model system for signal transduction. In haploid yeast cells, this pathway detects and transmits a pheromone signal emitted by cells of the opposite mating type to induce a mating response (*28*). As investment in mating carries high cost and leads to cell‐cycle arrest (*29*), induction of the pathway normally occurs with high precision (*30*) and overall low pathway noise (Figure S4; compare to Figure 1b).

**Figure 4.**
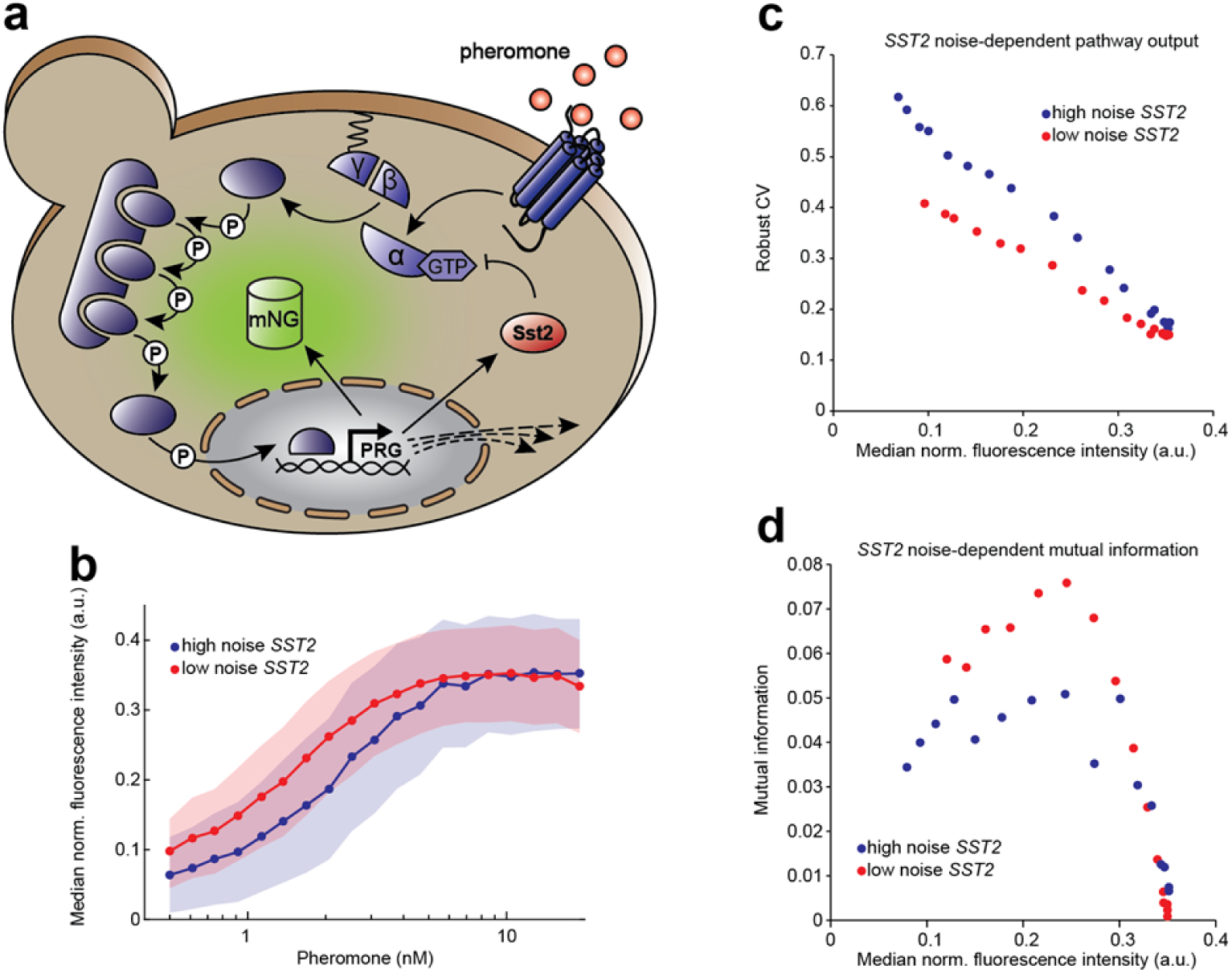
Calibration of noise in a signaling pathway component alters noise of pathway output. **(a) Schematic depiction of the yeast pheromone signaling (mating) pathway.** Yeast cells sense pheromones secreted by the opposite mating type via a G‐protein coupled receptor. The signal is transduced through a mitogen‐activated protein kinase (MAPK) cascade and ultimately induces the expression of pheromone response genes (PRG), such as the upstream negative feedback regulator Sst2. As a pathway activity readout, mNeongreen (mNG) driven by PRG promoter P_FUS1_ was genomically integrated. **(b) Dose response curves of the pathway reporter** Blue and red indicate high noise and low noise condition of *SST2* expression, respectively. Cells were stimulated for 180 minutes with different concentrations of pheromone. Lines indicate medians of normalized fluorescence, shaded areas show the corresponding median absolute deviations. Low noise (8.5 ng/µl doxycycline) and high noise (1 ng/µl doxycycline, 10 mM theophylline) conditions for *SST2* expression were chosen in order to achieve similar pathway responses. **(c) Pathway reporter noise at different pathway activity levels for high and low noise *SST2* conditions.** Points from left to right indicate stimulation with increasing pheromone concentrations as in **b**. Over the whole pathway activity range, the population of yeast cells with low noise in *SST2* expression displays a lower robust CV in the pathway reporter output than the population with high noise in *SST2* expression. **(d) Pathway information transmission is more precise with lower noise in *SST2* expression.** Mutual information between pheromone input and pathway output plotted against the pathway output for better comparison (see Methods for calculation). Precision is highest at intermediate reporter activity and overall higher with lower noise in Sst2. Points indicate stimulation with different pheromone concentrations as in **b** and **c**.

We applied the noise tuner to control expression of an upstream negative feedback regulator Sst2 that has been shown to act as noise suppressor of the pathway (*31*). Instead of measuring the regulated gene directly, in this case we quantified the pathway output by means of a mNeongreen reporter gene controlled by the promoter of the pathway response gene *FUS1*. For regulation of *SST2*, we tested different combinations of doxycycline and theophylline (Figure S5) and chose two that gave similar pathway output but were expected to result in different noise levels. Two populations with these different *SST2* expression settings were stimulated with different doses of pheromone, indeed resulting in similar pathway activity over the range of applied pheromone concentrations (Figure 4b). However, noise in the pathway output was substantially different over essentially the whole range of output levels (Figure 4c), with higher noise in Sst2 being unambiguously manifested in higher signaling and consequently output noise. From this we conclude that (i) Sst2 noise can be regulated by the noise tuner and (ii) changes in Sst2 noise are transmitted, at least qualitatively, through the cascade without being filtered out. We thus demonstrate that the noise tuner can be used for studying cellular networks, e.g. for identifying components whose noise‐characteristics are critical for robustness of the whole network.

Pathway noise impacts the amount of information that can be conveyed through the pathway. In order to quantify the precision at which cells could determine the pheromone concentration we calculated the mutual information between the input and output signals, which is a key metric for the accuracy of a signaling pathway (*32*). We found an up to 50% change in mutual information with low‐noise compared to high‐noise *SST2* settings (Figure 4d). Thus, noise changes generated by the noise tuner resulted in significant changes in signal transduction accuracy of the mating pathway.

## Conclusion

In this work, we present a novel approach to effectively decouple mean and noise of yeast gene expression, allowing us to flexibly adjust different levels for median expression and noise in clonally identical cells. The noise tuner acts on two basic rates of gene expression, with two small molecule inducers individually controlling transcription and mRNA degradation rates. In this system, the transcription rate determines the noise level, whereas adjustment of the mRNA degradation rate allows for similar mean expression levels with different promoter activities. We found that this is consistent with a mathematical model of gene expression, which shows that for genes with sufficiently high protein lifetime, transcript stability has only little effect on noise.

Comparison to constructs with native 3’‐RRs suggests that the low‐expression noise tuner operates at the lower end of the physiological mean‐noise space that can be conferred by 3’‐RRs. A more stable ribozyme shifts the expression towards higher levels while maintaining the noise tuning capabilities. The noise tuners presented here can produce more than twofold difference in noise over a fivefold range of mean expression levels. We demonstrate that the described noise control system could be used to study noise sensitivity of cellular networks and identifying pathway components which are critical for robustness (*33*).

Noise control by regulation of mRNA levels, as demonstrated here, may be not only highly effective but also metabolically cheap, given the generally low numbers of mRNA molecules (*34*), as compared to protein molecules, which are several magnitudes higher (*35*). Furthermore, due to its simplicity, the noise tuner might find common application in the design of synthetic genetic circuits that lack robustness if they do not contain noise dampening modules.

## Methods

### Plasmid and strains

All plasmids used in this study are listed in Table S1. Plasmids were either cut with PmeI (New England Biolabs) or PCR‐amplified to yield linear DNA with homology sequences on both ends. All primers are listed in Table S2. For PCR‐amplified DNA, the homology was introduced by primer overhangs of at least 50 bp. PCRs were performed using PrimeSTAR GXL DNA polymerase (Takara). The resulting single integration constructs were transformed into *Saccharomyces cerevisiae* strains using a standard lithium acetate protocol.

All *Saccharomyces cerevisiae* strains in this study derived from haploid SEY6210 mating type a and are listed in Table S3. The strain used for the proof of concept of direct noise control (YMFM050) was built by transformation of 1 μg of PmeI‐digested plasmid pMFM048. The plasmid contains an mNeongreen reporter gene, kind gift of Hyun Youk, driven by an inducible TetO7‐promoter. Downstream of the coding sequence the construct contains the L2b8‐a1‐t41 inducible ribozyme (*20*), provided by Christina Smolke) and the minimal terminator sequence T(Synth27) (*22*). To construct the plasmid, individual PCR fragments containing the parts were joined by Gibson assembly. The ribozyme was introduced using restriction ligation via AvrII and XhoI. Based on YMFM050, the strains with native terminator regions (YMFM101, 102, 103, 104) were constructed by *in vivo* tagging with PCR fragments created from primers with homology overhangs and the templates pAA263, pAA207, pAA208, pAA209, and pMFM065, respectively.

The *SST2* noise mating pathway reporter strain was constructed by *in vivo* tagging of strain YAA328 with a PCR fragment derived from pAA263 to integrate the TetO7 promoter upstream of the native *SST2* open reading frame and subsequently integrating a PCR fragment derived from pMFM058 to replace the native terminator region downstream of the *SST2* open reading frame with the inducible ribozyme and the minimal terminator.

### Media and growth conditions

All experiments were performed with cells grown in low fluorescent synthetic defined media (LD). Cells were inoculated from single colonies and grown at 30°C with shaking over night for at least 16 hours. Day cultures were inoculated to an initial OD of 0.05 and grown at 30°C with shaking for 5 to 6 hours to an OD between 0.4 and 0.8 prior to the measurement. For mating pathway stimulation experiments, media was supplied with 2 μM casein and cells were grown to at least OD 0.2 prior to stimulation with pheromone.

For strains harboring genes controlled by Tet‐promoter and inducible ribozyme, doxycycline and theophylline were added to the final concentrations as indicated in the results section. Due to the low solubility in aqueous solutions, LD containing theophylline was prepared by adding the appropriate amount of theophylline powder directly to the media.

1 L synthetic defined low fluorescent media contains 6.9 g YNB (Formedium), 790 mg of the appropriate amino acid mix (Formedium), and 2% glucose (Roth). Selection of transformants was done on agar plates containing YNB, glucose and the appropriate amino acid dropout mix, supplemented with 1.5 % agar (Becton Dickinson) for transformants with auxotrophic marker. Transformants with antibiotic marker KanMX were selected using plates containing YPD (Roth), 1.5 % agar and 500 mg/ mL G418 (Formedium).

### Flow cytometry measurements and noise calculation

Yeast cells were grown in 24‐well plates and then transferred to a 96‐well plate. Using a high throughput sampler, cells were injected into an LSR Fortessa Special Order flow cytometer (BD Biosciences). Fluorescent proteins were measured with different lasers, 488 nm for mNeongreen and 447 nm for the constitutively expressed mTurquoise2. Using the BD FACS DIVA software (BD Biosciences), cells were gated in a FSC‐A / SSC‐A plot to exclude debris. Per sample, 50,000 cells from within the gate were acquired.

To account for autofluorescence, a yeast strain was measured that contained the mTurquoise2 intrinsic control module but not the mNeongreen gene. The signal for green fluorescence of every individual cell was compared to the distribution of the corresponding signals of the autofluorescence strain. Treating the background distribution as a probability distribution, a random value of that distribution (based on the probability) was subtracted from the measured fluorescence of each individual cell. When the value was within the range of the background distribution, the random value was picked only from the part of the distribution that was lower than the measured value. This method of background subtraction works under the prerequisite that the background fluorescence of a given cell is bounded by 0 and the fluorescence signal of that cell. As a result, for distributions that overlap with the distribution of the autofluorescence control, cells towards the left end of a population are deducted a smaller value than cells towards the right end of the population.

The median absolute deviation is given as

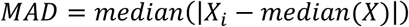

with the fluorescence intensity of an individual cell, *X_i_*. Median and MAD were calculated for the set of autofluorescence‐subtracted values and cells that deviated from the median by more than five times the scaled MAD (=MAD/0.6745) were excluded from the subsequent analysis. The resulting value for each cell was then divided by the fluorescence value of the constitutively expressed mTurquoise2 expression control to normalize for the general expression state of each individual cell. Median and MAD were calculated from the resulting set of values and outliers were removed as above. A final median and MAD was calculated for the remaining cells. As a measure for noise, the robust coefficient of variation (rCV) was calculated as

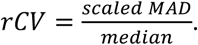

Iso‐median lines (Figures 2a, 3b, S3) were plotted using MATLAB’s *contour* function.

### Stochastic simulation

The simulation is based on a two‐stage model of gene expression as has been analyzed by Shahrezaei and Swain (*25*). The model describes the probability distribution of protein numbers in a cell, based on the underlying probabilities of mRNA synthesis *v*_0_, mRNA degradation *d*_0_, protein synthesis *v*_1_ and protein degradation (*d*_1_, see also Figure 3a). Given the assumption that the protein lifetime is much longer than the lifetime of a transcript 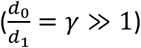, a probability distribution can be derived for the steady state protein number *n*:

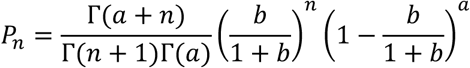

where the production and degradation probabilities are reformulated as mRNA production per protein lifetime 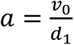 and the translational burst size 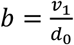 Г() denotes the gamma function to take non‐integer values into account.

To confirm that the CV becomes independent of the translational burst size *b* as long as *b* ≫ 1, we performed stochastic simulations of the four processes above using the Gillespie formulism (*36*) as was also done by Shahrezaei and Swain. We fixed the protein half‐life to be two hours, which is a conservative estimate based on of GFP data (*37*), resulting in the protein degradation rate 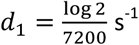. We varied the parameter *a*, thereby achieving a corresponding range in values of the mRNA synthesis rate *v*_0_. Up to a normalization, the set of values for *a* and *b* were chosen by non‐negative matrix factorization (using the function *nnmf* in MATLAB) of the matrix of experimental mean fluorescent values (similar to Figure 2a). Since the mean is given theoretically by *ab*, this provides a conversion from the doxycycline and theophylline concentrations used in the experiment to (up to a factor) the corresponding *a* and *b* values. The normalization factors were chosen to match the experimental range of rCV and to give reasonable values for the mean number of molecules and the burst size. Finally, the protein synthesis rate was fixed *v*_1_ *=* 0.11 s^‐1^ such that the range of mRNA half‐lives fell within the biologically expected range (minutes).

The resultant values were

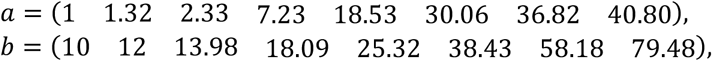

which corresponded to mRNA synthesis rates (in units of hour^‐1^) of

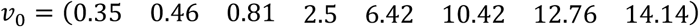

and mRNA half‐lives (in units of mins) of

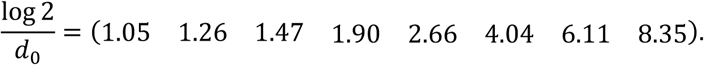

Simulations were performed using these values and the properties of the steady‐state protein distribution were compared to the experimental result (Figure 2a and 3b).

### Calculation of mutual information

Mutual information describes how much information (in bits) from an input distribution (here: pheromone concentrations) can be inferred from an output distribution (here: fluorescence intensities). In the experiments described in this study, only the output distribution is known (continuous data sets of fluorescence intensities), whereas the input can only be described by discrete values (20 pheromone concentrations ranging from 0 nM to 18.75 nM). We used the formula derived in (*38*) to calculate the mutual information *I* between a discrete dataset *X* and a continuous dataset *Y*:

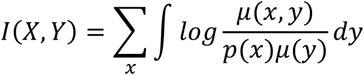

with the discrete probability function *p(x)* and the continuous densities *µ(x, y)* and *µ(y)*. Based on (*39*) we calculated *I(X, Y)* for a moving window of four experimental subsets of subsequent pheromone concentrations, considering pheromone/ fluorescence data of 45,000 cells for each subset. For the 20 pheromone concentrations tested, this resulted in 17 values for mutual information (first window: 1 to 4; last window: 17 to 20).

We note that the absolute values of mutual information calculated is somewhat arbitrary as it depends heavily on the number of different inputs measured: A high‐resolution dose‐response with many input pheromone concentrations contains only little mutual information when adjacent concentrations are compared. In contrast, when only off‐ and on‐state of the pathway are compared (0 nM and 18.75 nM pheromone), the mutual information increases but the results would not allow to quantify differences between the high‐noise and the low‐noise setting.

## References

1. M. B. Elowitz, A. j. Levine, E. D. Siggia, P. S. Swain, Stochastic Gene Expression in a Single Cell. Science. 297, 1183–1186 (2002), doi:10.1126/science.1070919.

2. P. Shah, Y. Ding, M. Niemczyk, G. Kudla, J. B. Plotkin, Rate-limiting steps in yeast protein translation. Cell. 153, 1589–1601 (2013), doi:10.1016/j.cell.2013.05.049.

3. G. Chalancon et al., Interplay between gene expression noise and regulatory network architecture. Trends in genetics: TIG. 28, 221–232 (2012), doi:10.1016/j.tig.2012.01.006.

4. Newman, John R. S. et al., Single-cell proteomic analysis of S. cerevisiae reveals the architecture of biological noise. Nature. 441, 840–846 (2006), doi:10.1038/nature04785.

5. Z. Wang, J. Zhang, Impact of gene expression noise on organismal fitness and the efficacy of natural selection. Proceedings of the National Academy of Sciences. 108, E67–E76 (2011), doi:10.1073/pnas.1100059108.

6. B. Lehner, Selection to minimise noise in living systems and its implications for the evolution of gene expression. Mol Syst Biol. 4, 1389 (2008), doi:10.1038/msb.2008.11.

7. A. Eldar, M. B. Elowitz, Functional roles for noise in genetic circuits. Nature. 467, 167–173 (2010), doi:10.1038/nature09326.

8. L. Keren et al., Massively Parallel Interrogation of the Effects of Gene Expression Levels on Fitness. Cell. 166, 1282–1294.e18 (2016), doi:10.1016/j.cell.2016.07.024.

9. S. Hooshangi, S. Thiberge, R. Weiss, Ultrasensitivity and noise propagation in a synthetic transcriptional cascade. Proceedings of the National Academy of Sciences of the United States of America. 102, 3581–3586 (2005), doi:10.1073/pnas.0408507102.

10. A. Becskei, L. Serrano, Engineering stability in gene networks by autoregulation. Nature. 405, 590–593 (2000).

11. I. Lestas, G. Vinnicombe, J. Paulsson, Fundamental limits on the suppression of molecular fluctuations. Nature. 467, 174–178 (2010), doi:10.1038/nature09333.

12. K. F. Murphy, R. M. Adams, X. Wang, G. Balázsi, J. J. Collins, Tuning and controlling gene expression noise in synthetic gene networks. Nucleic Acids Research. 38, 2712–2726 (2010), doi:10.1093/nar/gkq091.

13. A. Aranda-Díaz, K. Mace, I. Zuleta, P. Harrigan, H. El-Samad, Robust Synthetic Circuits for Two-Dimensional Control of Gene Expression in Yeast. ACS synthetic biology. 6, 545–554 (2017), doi:10.1021/acssynbio.6b00251.

14. J. M. Raser, E. K. O’Shea, Control of Stochasticity in Eukaryotic Gene Expression. Science. 304, 1811–1814 (2004), doi:10.1126/science.1098641.

15. W. J. Blake, M. Kaern, C. R. Cantor, J. J. Collins, Noise in eukaryptic gene expression. Nature. 422, 633–637 (2003).

16. M. Thattai, A. van Oudenaarden, Intrinsic noise in gene regulatory networks. PNAS. 98, 8614–8619 (2001).

17. K. E. Galloway, E. Franco, C. D. Smolke, Dynamically Reshaping Signaling Networks to Program Cell Fate via Genetic Controllers. Science. 341, 12350051–123500511 (2013), doi:10.1126/science.1235005.

18. T. C. Williams, L. K. Nielsen, C. E. Vickers, Engineered Quorum Sensing Using Pheromone-Mediated Cell-to-Cell Communication in Saccharomyces cerevisiae. ACS Synth. Biol. 2, 136–149 (2012), doi:10.1021/sb300110b.

19. H. Youk, W. A. Lim, Secreting and Sensing the Same Molecule Allows Cells to Achieve Versatile Social Behaviors. Science. 343, 1242782 (2014), doi:10.1126/science.1242782.

20. J. C. Liang, A. L. Chang, A. B. Kennedy, C. D. Smolke, A high-throughput, quantitative cell-based screen for efficient tailoring of RNA device activity. Nucleic Acids Research. 40, e154 (2012), doi:10.1093/nar/gks636.

21. S. Urlinger et al., Exploring the sequence space for tetracycline-dependent transcriptional activators: Novel mutations yield expanded range and sensitivity. PNAS. 97, 7963–7968 (2000).

22. K. A. Curran et al., Short Synthetic Terminators for Improved Heterologous Gene Expression in Yeast. ACS synthetic biology. 4, 824–832 (2015), doi:10.1021/sb5003357.

23. N. C. Shaner et al., A bright monomeric green fluorescent protein derived from Branchiostoma lanceolatum. Nature methods. 10, 407–409 (2013), doi:10.1038/nmeth.2413.

24. M. Yamanishi et al., A Genome-Wide Activity Assessment of Terminator Regions in Saccharomyces cerevisiae Provides a ″Terminatome″ Toolbox. ACS Synth. Biol. 2, 337–347 (2013), doi:10.1021/sb300116y.

25. V. Shahrezaei, P. S. Swain, Analytical distributions for stochastic gene expression. PNAS. 105, 17256–17261 (2008).

26. A. Raj, A. van Oudenaarden, Single-molecule approaches to stochastic gene expression. Annual review of biophysics. 38, 255–270 (2009), doi:10.1146/annurev.biophys.37.032807.125928.

27. D. R. Larson, R. H. Singer, D. Zenklusen, A single molecule view of gene expression. Trends in Cell Biology. 19, 630–637 (2009), doi:10.1016/j.tcb.2009.08.008.

28. C. G. Alvaro, J. Thorner, Heterotrimeric G Protein-coupled Receptor Signaling in Yeast Mating Pheromone Response. The Journal of biological chemistry. 291, 7788–7795 (2016), doi:10.1074/jbc.R116.714980.

29. A. Banderas, M. Koltai, A. Anders, V. Sourjik, Sensory input attenuation allows predictive sexual response in yeast. Nat Comms. 7, 12590 (2016), doi:10.1038/ncomms12590.

30. M. K. Malleshaiah, V. Shahrezaei, P. S. Swain, S. W. Michnick, The scaffold protein Ste5 directly controls a switch-like mating decision in yeast. Nature. 465, 101–105 (2010), doi:10.1038/nature08946.

31. G. Dixit, J. B. Kelley, J. R. Houser, T. C. Elston, H. G. Dohlman, Cellular noise suppression by the regulator of G protein signaling Sst2. Molecular cell. 55, 85–96 (2014), doi:10.1016/j.molcel.2014.05.019.

32. C. Waltermann, E. Klipp, Signal integration in budding yeast. Biochemical Society transactions. 38, 1257–1264 (2010), doi:10.1042/BST0381257.

33. J. Masel, M. L. Siegal, Robustness. Trends in genetics: TIG. 25, 395–403 (2009), doi:10.1016/j.tig.2009.07.005.

34. M. J. Holland, Transcript abundance in yeast varies over six orders of magnitude. The Journal of biological chemistry. 277, 14363–14366 (2002), doi:10.1074/jbc.C200101200.

35. S. Ghaemmaghami et al., Global analysis of protein expression in yeast. Nature. 425, 737–741 (2003).

36. D. T. Gillespie, Exact stochastic simulation of coupled chemical reactions. J. Phys. Chem. 81, 2340–2361 (1977), doi:10.1021/j100540a008.

37. A. Natarajan, S. Subramanian, F. Srienc, Comparison of mutant forms of the green fluorescent protein as expression markers in Chinese hamster ovary (CHO) and Saccharomyces cerevisiae cells. Journal of Biotechnology. 62, 29–45 (1998), doi:10.1016/S0168-1656(98)00040-6.

38. B. C. Ross, Mutual information between discrete and continuous data sets. PLoS ONE. 9, e87357 (2014), doi:10.1371/journal.pone.0087357.

39. A. Borst, F. E. Theunissen, Information theory and neural coding. Nature neuroscience. 2, 947–957 (1999), doi:10.1038/14731.

